# Ventral Tegmental Area Glutamate Neurons establish a mu-opioid receptor gated circuit to mesolimbic dopamine neurons and regulate prescription opioid-seeking behavior

**DOI:** 10.1101/2022.12.11.519972

**Authors:** Dillon J. McGovern, Abigail M. Polter, Annie Ly, Emily D. Prevost, Connor McNulty, David H. Root

## Abstract

A two-neuron model of ventral tegmental area (VTA) opioid function classically involves VTA GABA neuron regulation of VTA dopamine neurons via a mu-opioid receptor dependent inhibitory circuit. However, this model predates the discovery of a third major type of neuron in the VTA: glutamatergic neurons. We find that about one-quarter of VTA neurons expressing the mu-opioid receptor are glutamate neurons without molecular markers of GABA co-release. Glutamate-Mu opioid receptor neurons are topographically distributed in the anterior VTA. The majority of remaining VTA mu-opioid receptor neurons are GABAergic neurons that are largely within the posterior VTA and do not express molecular markers of glutamate co-release. Optogenetic stimulation of VTA glutamate neurons results in monosynaptic excitatory currents recorded from VTA dopamine neurons that are reduced by presynaptic activation of the mu-opioid receptor *ex vivo*, establishing a local mu-opioid receptor dependent excitatory circuit from VTA glutamate neurons to VTA dopamine neurons. This VTA glutamate to VTA dopamine pathway regulates dopamine release to the nucleus accumbens through mu-opioid receptor activity *in vivo*. Behaviorally, VTA glutamate calcium-related neuronal activity increased following oxycodone consumption and response-contingent oxycodone-associated cues during self-administration and abstinent reinstatement of drug-seeking behavior. Further, chemogenetic inhibition of VTA glutamate neurons reduced abstinent oxycodone-seeking behavior in male but not female mice. These results establish 1) a three-neuron model of VTA opioid function involving a mu-opioid receptor gated VTA glutamate neuron pathway to VTA dopamine neurons that controls dopamine release within the nucleus accumbens, and 2) that VTA glutamate neurons participate in prescription opioid-seeking behavior.

## INTRODUCTION

The United States continues to struggle with an opioid epidemic involving unprecedented numbers of opioid-related deaths[1]. Both prescription and synthetic opioid-related overdoses have been exacerbated by the COVID19 pandemic[2]. Understanding the neuronal cell-types and circuits affected by opioids may provide new insights into mitigating the severity of the opioid epidemic. Heroin, morphine, fentanyl, oxycodone, and other addiction-related opioids bind to the µ-opioid receptor (MOR) in the central nervous system[3]. Genetic ablation of MOR in mice results in resistance to the rewarding effects of morphine as well as naloxone-precipitated withdrawal[4], indicating that the MOR significantly contributes to rewarding and aversive components of opioid addiction. While neurons expressing the MOR are located throughout the brain, MOR binding within the ventral tegmental area (VTA) is necessary and sufficient for opioid reward. In rodents, activation of the MOR in the VTA is highly rewarding[5] and blocking the MOR in the VTA prevents the reinforcing aspects of opioids[6] as well as the self-administration of opioids[7]. Thus, µ-opioid receptors within the VTA are thought to play a critical role opioid addiction.

Thirty years ago, a two-neuron model of VTA opioid function was proposed[8]. At this time, two types of VTA neurons were known: neurons that release the neuromodulator dopamine and neurons that release the inhibitory neurotransmitter GABA. Electrophysiological studies of VTA neurons suggested that VTA GABA neurons express the MOR and directly regulate VTA dopamine neurons via presynaptic actions[8].The two-neuron model of opioid addiction has been highly replicated to show that MOR activation: 1) inhibits VTA GABA neurons and terminals resulting in the disinhibition of VTA dopamine neurons to cause opioid reward, and 2) blockade of the MOR in opioid-dependent subjects activates VTA GABA neurons to engender the aversive effects of opioid withdrawal[9-13].

While highly influential, recent investigations suggest a need to revise the two-neuron model of VTA opioid function. Electrophysiological recording studies have shown that only a subset of VTA GABA neurons show MOR-elicited electrophysiological responses suggestive of MOR-expression[14,15]. Additional work heavily implicates rostromedial tegmental area GABA neurons in opioid signaling and behavior[16]. A subset of VTA glutamate neurons, discovered fifteen years after the development of the two-neuron model of VTA opioid function[17], expresses MOR mRNA and shows MOR-elicited electrophysiological responses supportive of MOR-expression[18-20]. Further, VTA glutamate receptor activation is required for the increased firing of dopamine neurons in response to morphine[21] and heroin self-administration[22]. These findings indicate that current understanding of the VTA cell-types and circuits regulated by the MOR are not wholly understood.

We find that VTA MOR-expressing cell-types are largely divided between GABA neurons defined by the vesicular GABA transporter (VGaT) and glutamate neurons defined by the vesicular glutamate transporter 2 (VGluT2). VTA VGaT+MOR+ neurons predominate the posterior VTA while the majority of anterior VTA MOR-expressing neurons are VGluT2 +MOR+ neurons. Both MOR cell-types largely lack markers of glutamate and GABA co-transmission (VGluT2+VGaT+), suggesting that each VTA MOR subpopulation is a subtype of VTA VGluT2―VGaT+ or VGluT2+VGaT― neurons. Further, we identified a local circuit from VTA VGluT2 neurons to VTA dopamine neurons that is regulated by the MOR using *ex vivo* slice electrophysiology. *In vivo*, this MOR-regulated VTA VGluT2 to VTA dopamine circuit modulates the release of dopamine in the nucleus accumbens. Finally, we show that VTA VGluT2 neurons are activated by and in males, necessary for cued reinstatement of prescription opioid-seeking behavior. Together, we identify an unexpected MOR-regulated mesolimbic circuitry from VTA VGluT2+ MOR+ neurons that participates in prescription opioid-seeking behavior.

## RESULTS

### Cell-type specific expression of VTA MOR and topographic distribution

Given that expression of the MOR by VTA GABA neurons is well-established, and that VTA VGluT2+ VGaT+ neurons are a subset of VTA GABA neurons [28], we determined which cell-types express the MOR across the VTA. Thin sections of the VTA from wildtype C57 mice were examined for the expression of transcripts encoding *VGluT2* (for the detection of VTA glutamate-releasing neurons), *VGaT* (for the detection of VTA GABA-releasing neurons), and *Oprm1* (for the detection of MOR-expressing neurons) (**Figure 1A**). To determine a topographical distribution of the MOR on VTA VGluT2+ and VGaT+ neurons, VTA sections prior to the full emergence of the interpeduncular nucleus were designated as the anterior VTA while those afterwards were designated as the posterior VTA. In the anterior VTA, 56% ± 8.23% of MOR-neurons expressed VGluT2 without VGaT, 29% ± 8.46% expressed VGaT without VGluT2, and 11% ± 2.26% triple expressed the MOR with VGluT2 and VGaT transcripts (**Figure 1B**). In the posterior VTA, the distribution of MOR expressing cell-types shifted, with a predominance of MOR-neurons expressing VGaT without VGluT2 (88% ± 3.5%), and only 9% ± 2.74% expressing VGluT2 without VGaT, and 1.86% ± 0.30% expressing both VGluT2 and VGaT (**Figure 1C**).

**Figure 1:**
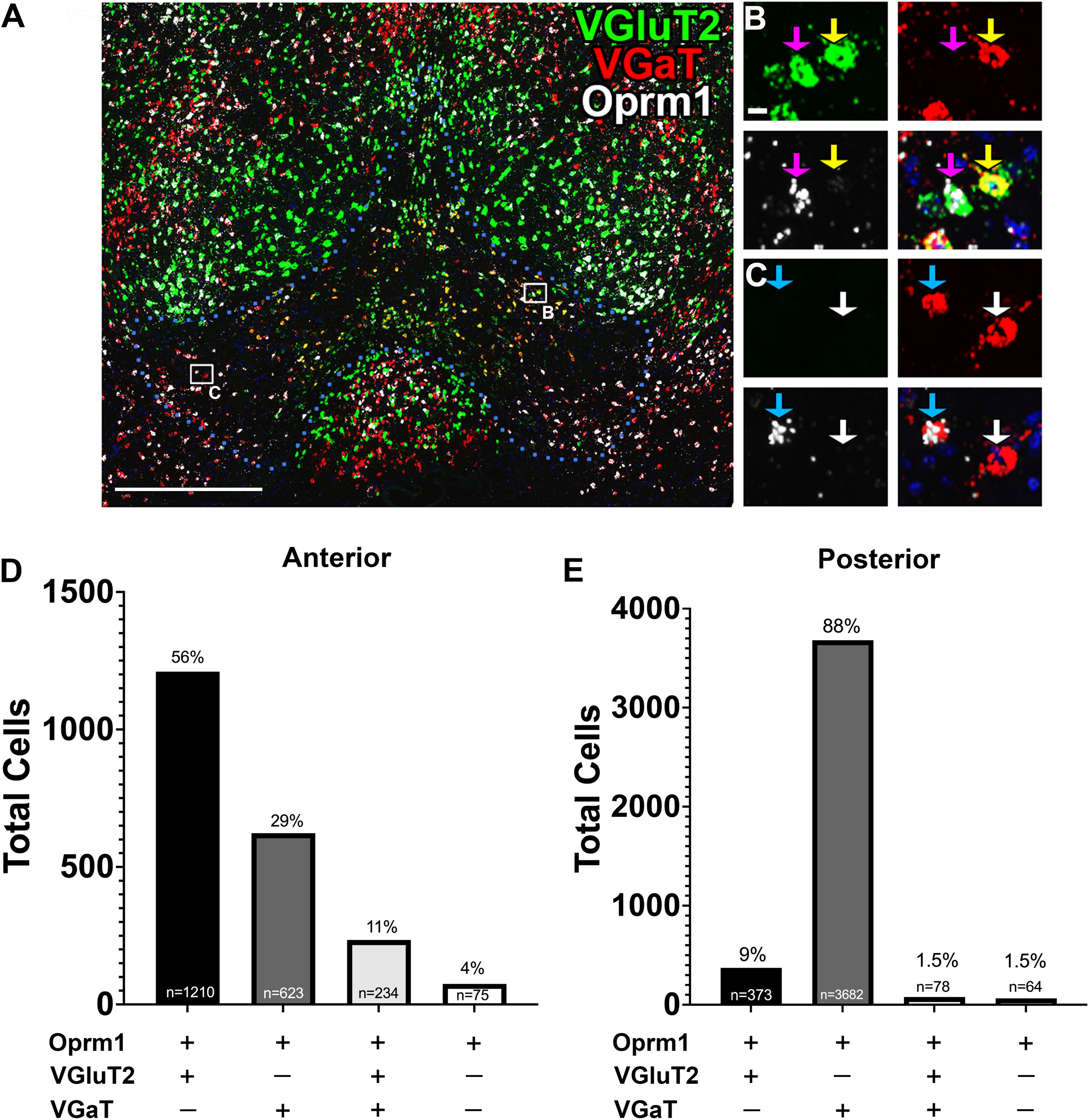
VTA VGluT2+ mu-opioid receptor distribution. **A**. VTA section labeled for RNA transcripts encoding VGluT2 (green), VGaT (red), oprm1 (white) mRNAs. Dotted blue represents VTA border. **B**. Example VTA neuron co-expressing VGluT2 and Oprm1 without VGaT mRNA (purple arrow) as well as a VGluT2 and VGaT co-expressing neuron lacking Oprm1 mRNA (yellow arrow). **C**. Example VTA neurons that co-express VGaT and Oprm1 mRNA (blue arrow) or express VGaT without Oprm1 mRNA. **D**. Anterior distribution (AP: 3.0-3.4 from bregma), highest percentage of Oprm1 mRNA neurons co-expressed VGluT2 only (56% ± 8.23%), followed by Oprm1 co-expression with VGaT only (29% ± 8.46%), and Oprm1 co-expression with VGluT2 and VGaT (11% ± 2.26%). Cell counts anterior: (VGluT2+VGaT―: 172.85 ± 17.86, VGluT2―VGaT+: 89 ± 29.99, VGluT2+VGaT+: 33.43 ± 6.51). **E**. Posterior distribution (AP: 3.4-3.8 from bregma), highest percentage of Oprm1 mRNA neurons co-expressed with VGaT only (88% ± 3.5%), followed by Oprm1 co-expression with VGluT2 only (9% ± 2.74%), and VGluT2+VGaT+ co-expression (1.86% ± 0.30%). Cell counts posterior (VGluT2+VGaT―: 53.29 ± 9.71, VGluT2―VGaT+: 526 ± 91.88, VGluT2+VGaT+: 11.14 ± 1.90). All values represent the mean ± SEM. Scale bar in A is 500 µm and scale bar in B is 10 µm and applies to B-C.

Together, in anterior VTA, MOR is mostly expressed in VGluT2+VGaT― neurons and in posterior VTA, MOR is predominantly expressed in VGluT2― VGaT+ neurons. Thus, VGluT2+MOR+ neurons and VGaT +MOR+ neurons are genetically distinct subsets of VGluT2+VGaT― or VGluT2―VGaT+ neurons, respectively. There were no significant differences between male and female VTA cell-type distributions (**Supplemental Figure 1**).

### VTA VGluT2 neuron synapses onto local VTA dopamine neurons are regulated by the MOR

Based on the well-established MOR-regulated VTA GABA to VTA dopamine neuron microcircuit[8], we hypothesized that VTA VGluT2+MOR+ neurons establish MOR-sensitive local glutamatergic synapses onto VTA dopamine neurons. To test this hypothesis, we expressed CoChR tethered to GFP in VTA VGluT2 neurons and recorded from nonGFP VTA neurons *ex vivo* under monosynaptic testing conditions (TTX and 4-AP; **Figure 2A**). To determine the identity of the recorded postsynaptic neurons, we examined whether biocytin-filled neurons co-expressed tyrosine hydroxylase (TH) immunoreactivity. Of twelve sections examined, we recovered the biocytin-expressing neuron in five sections and all co-expressed tyrosine hydroxylase (TH) immunoreactivity (**Figure 2B**). Consistent with prior reports[29,30], we found that optical stimulation of VTA VGluT2-CoChR neurons resulted in mono-synaptic EPSCs on local VTA neurons (**Figure 2C**). Application of the MOR-selective agonist DAMGO (500 nM) significantly reduced currents elicited by activation of VTA VGluT2-CoChR neurons (Baseline amplitude= 49.71 ± 8.6 pA, DAMGO amplitude = 31.08 ± 7.1 pA, p = 0.014) (**Figure 2D**). In all neurons that showed a DAMGO-induced reduction in VGluT2-CoChR currents, the paired-pulse ratio significantly increased following DAMGO application (Baseline PPR = 0.87 ± 0.1, DAMGO PPR = 1.18 ± 0.1, p = 0.003) (**Figure 2E**), consistent with a presynaptic site of MOR activation. Together these results indicate that VTA VGluT2+MOR+ neurons establish monosynaptic MOR-regulated local glutamatergic synapses onto VTA dopamine neurons (**Figure 2F**).

**Figure 2.**
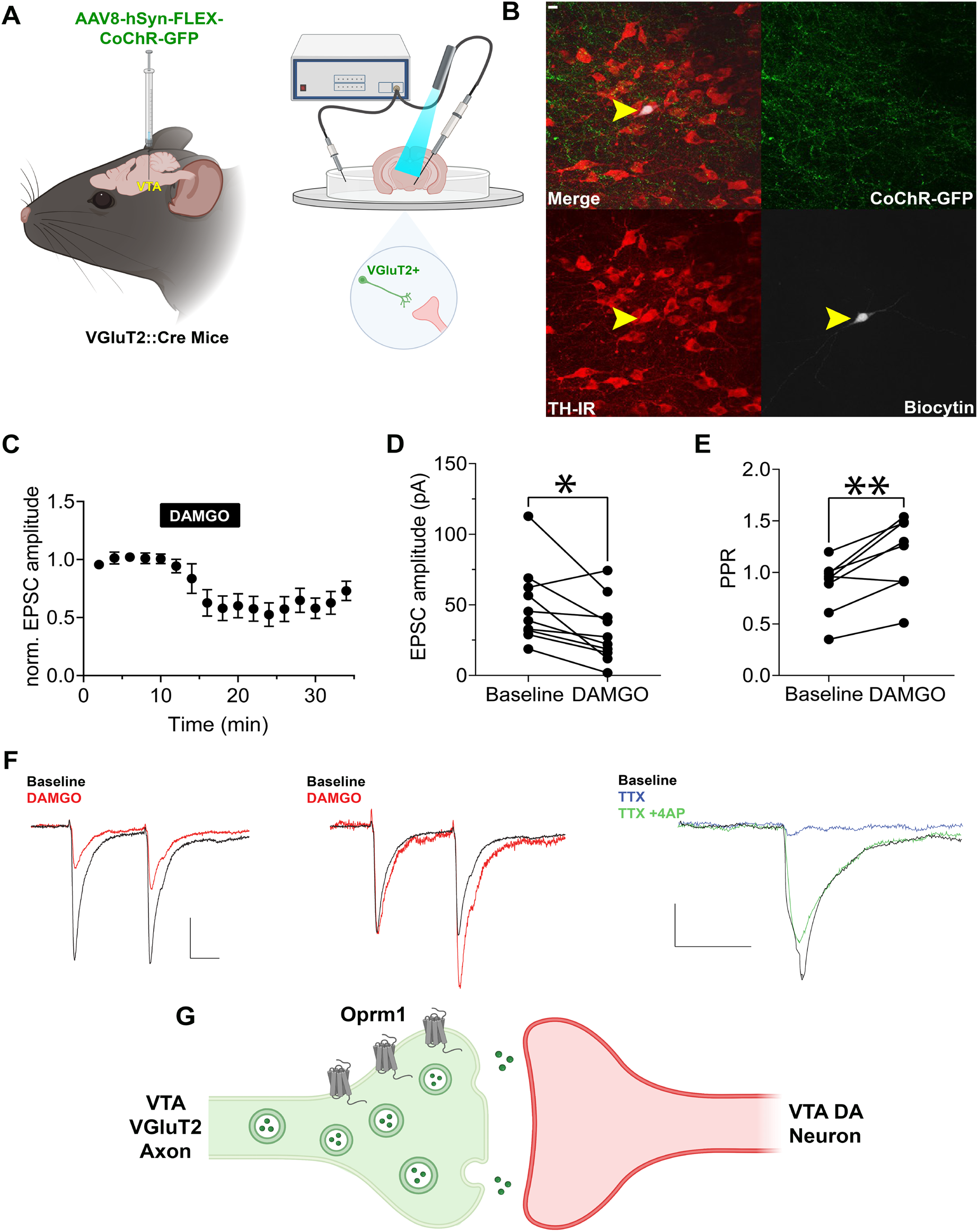
VTA VGluT2+ neurons establish a mu-opioid receptor regulated microcircuit to VTA dopamine neurons. **A**. Experimental schematic: VGluT2-IRES::Cre mice received a Cre-dependent injection encoding CoChR-GFP in the medial VTA. Brains were extracted and slices were prepared for whole cell recording. **B**. Immunohistochemical validation of local circuit, VTA VGluT2-CoChR-GFP processes (green) proximal to biocytin-filled neuron (white) co-labeled with tyrosine hydroxylase (red). **C**. Normalized EPSCs following VTA VGluT2 neuron optical stimulation. DAMGO wash at 10 minutes reduced optically-elicited EPSCs. **D**. Recorded neurons significantly reduced EPSC in recorded neurons (n=9/10), p=0.014. **E**. Significant increase in paired pulse ratio (PPR), p=0.003, suggesting pre-synaptic mu-opioid receptor regulation. **F**. Left: Representative 20ms trace, baseline (black), DAMGO (red); Middle: Scaled 20 ms trace, demonstrating PPR change; Right: Representative 20 ms trace, baseline (black), TTX (blue), TTX + 4AP (green). **G**. Proposed local circuit based on electrophysiological recording data: VTA VGluT2+MOR+ neurons monosynaptically cause EPSCs in VTA TH+ neurons, which is reduced by activation of the mu-opioid receptor on VTA VGluT2-neuron presynaptic terminals. Scale bar, upper left: 10 µm, Scale bar for representative EPSC traces: 20 pA, 20ms

### Nucleus accumbens dopamine release is regulated by a MOR-gated VTA VGluT2 neuron projection to VTA dopamine neurons

Having established a VTA VGluT2+MOR+ neuron microcircuit to VTA dopamine neurons, we aimed to test the hypothesis that this circuit influences dopamine release within the nucleus accumbens *in vivo*. To test this hypothesis, we expressed the red light-shifted ion channel ChRmine[31] in VTA VGluT2 neurons and in the same animals expressed the green light sensitive optical dopamine sensor GRABDA1h[32] in nucleus accumbens core (**Figure 3A; Supplemental Figure 2**). We then stimulated VTA VGluT2 neurons and recorded accumbal GRABDA activity *in vivo* following saline or oxycodone injection. ChRmine activation of VTA VGluT2 neurons resulted in increased GRABDA signal in nucleus accumbens core. While baseline GRABDA activity did not change between saline and oxycodone treatments (F(2,12) = 3.075, p = 0.084), accumbal GRABDA activity significantly differed between saline and oxycodone treatments following ChRmine stimulation of VTA VGluT2 neurons (F(2,12) = 5.168, p = 0.024). Posthoc contrast tests showed that oxycodone administration significantly reduced GRABDA activity following ChRmine activation of VTA VGluT2 neurons compared with each saline treatment (saline before oxycodone: F(1,6) = 10.412, p = 0.018; saline after oxycodone: F(1,6) = 8.847, p = 0.025) (**Figure 3B-E**). Together, accumbens core dopamine release elicited by VTA VGluT2 neuron stimulation that activates VTA dopamine neurons, is regulated by the MOR *in vivo* (**Figure 3F**).

**Figure 3.**
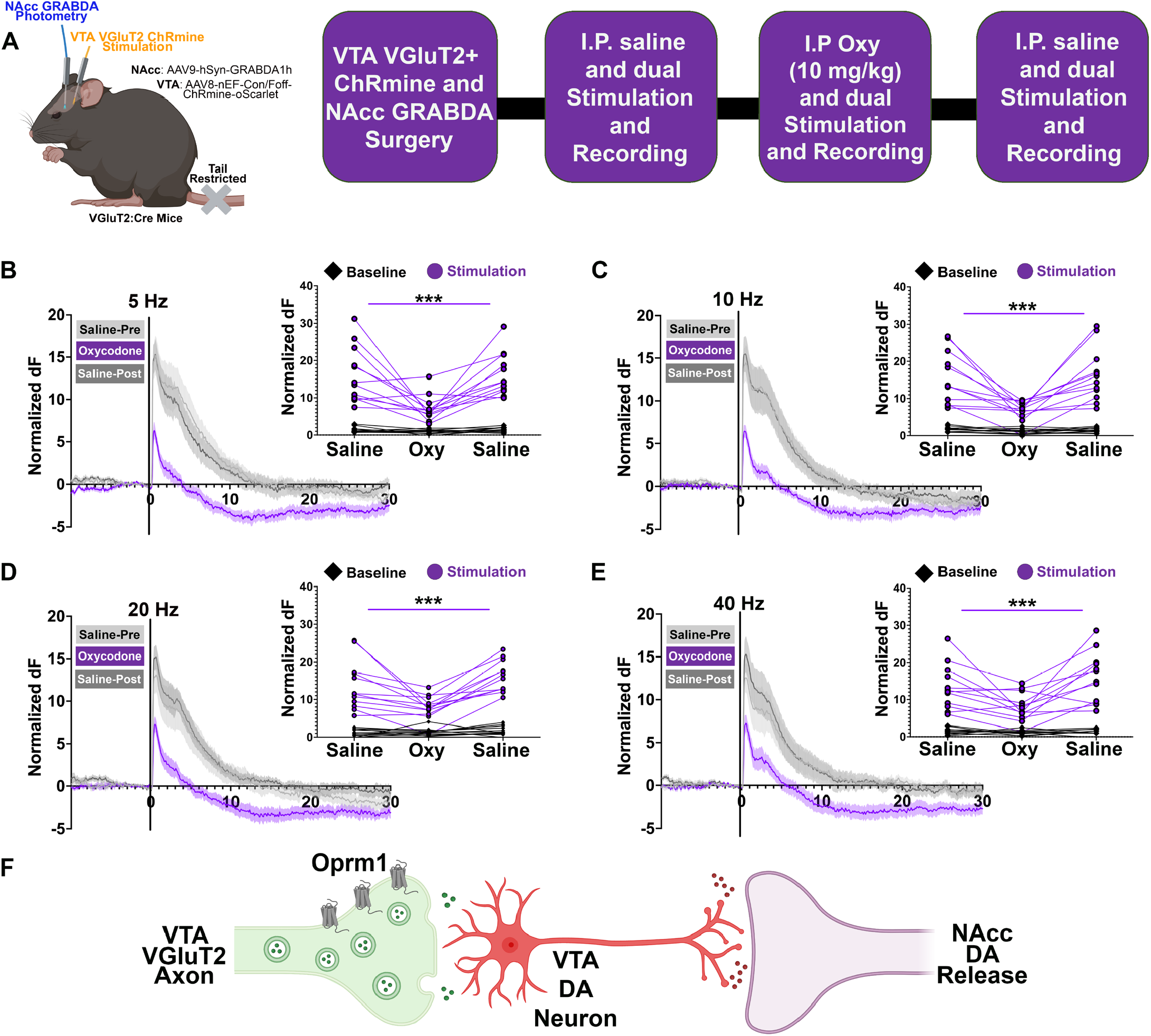
Dopamine release in the NAcc elicited by VTA VGluT2 neurons is reduced following oxycodone administration. **A-E**. Surgical schematic: AAV-GRABDA was injected in the nucleus accumbens core and implanted with a recording fiber optic. Medial VTA was injected with AAV-FLEX-ChRmine and implanted with a stimulating optic fiber. Experimental timeline: Four weeks following injections and implantations, mice were tail-restricted and administered an IP injection of saline. Five optical trains of 589nm light were delivered to VTA VGluT2 neurons at 5 Hz (**B**), 10 Hz (**C**), 20 Hz (**D**), 40 Hz (**E**) (order counterbalanced). Three days following saline pre-test, mice were administered an IP injection of oxycodone (10 mg/kg) and stimulated under the same conditions. Three days later, mice were given an IP saline injection and VTA VGluT2 neurons were stimulated once more. **F**. Proposed circuit: activated mu-opioid receptors present on VTA VGluT2 neuron terminals within the VTA reduce excitation onto local VTA neurons that release dopamine in the nucleus accumbens core.

### VTA VGluT2 neurons are activated by opioid-seeking behavior

After establishing a MOR-mediated VTA VGluT2 to VTA dopamine neuron pathway regulating accumbal dopamine release, we aimed to test the involvement of VTA VGluT2 neurons in opioid-seeking behavior. To accomplish this, we expressed the calcium-sensor GCaMP6m in VTA VGluT2+ neurons and recorded population changes in intracellular calcium during oxycodone self-administration or cue-induced reinstatement of extinguished drug-seeking behavior (**Figure 4A; Supplemental Figure 3**). Mice trained on a fixed ratio 2 reinforcement schedule to operantly self-administer oral oxycodone. A tone cue was paired with drug delivery to establish a drug-cue association (**Figure 4B**). Due to sex differences in self-administration of oral oxycodone[26] we separated female and male mice to evaluate sex as a biological variable. During self-administration, VTA VGluT2+ GCaMP activity significantly increased following response-contingent presentation of the oxycodone-associated cues, F(1,14)=22.87, p=0.0003, for female mice: t(8)=3.615, p=0.0056, and male mice: t(8) = 3.148, p=0.0142. Area under the curve (AUC) also significantly increased for oxycodone active poke F(1,14)=15.41, p=0.0015, for female mice: t(8)=2.860, p=0.0250, and male mice: t(8)=2.691, p=0.0348. No significant sex differences were observed between 5 maximum response F(1,14)=0.0023, p=0.9622 or area under the curve F(1,14)= 0.01433, p=0.9064. VTA VGluT2+ GCaMP activity significantly increased following head entry into the oxycodone magazine where drug consumption occurred, F(1,14)=40.21, p < 0.0001, for female mice: t(8)=5.580, p=0.0001, and male mice: t(8)=3.562 p=0.0062. AUC was also significantly different from baseline for oxycodone magazine entry, F(1,14)=36.83, p<0.0001, for female mice: t(8)=5.120, p=0.0003, and male mice: t(8)=3.462, p=0.0076 mice. There were no significant sex differences between maximum recorded fluorescence F(1,14)=0.344, p=0.566 or area under the curve, F(1,14)=1.402, p=0.256 (**Figure 4C-H**).

**Figure 4.**
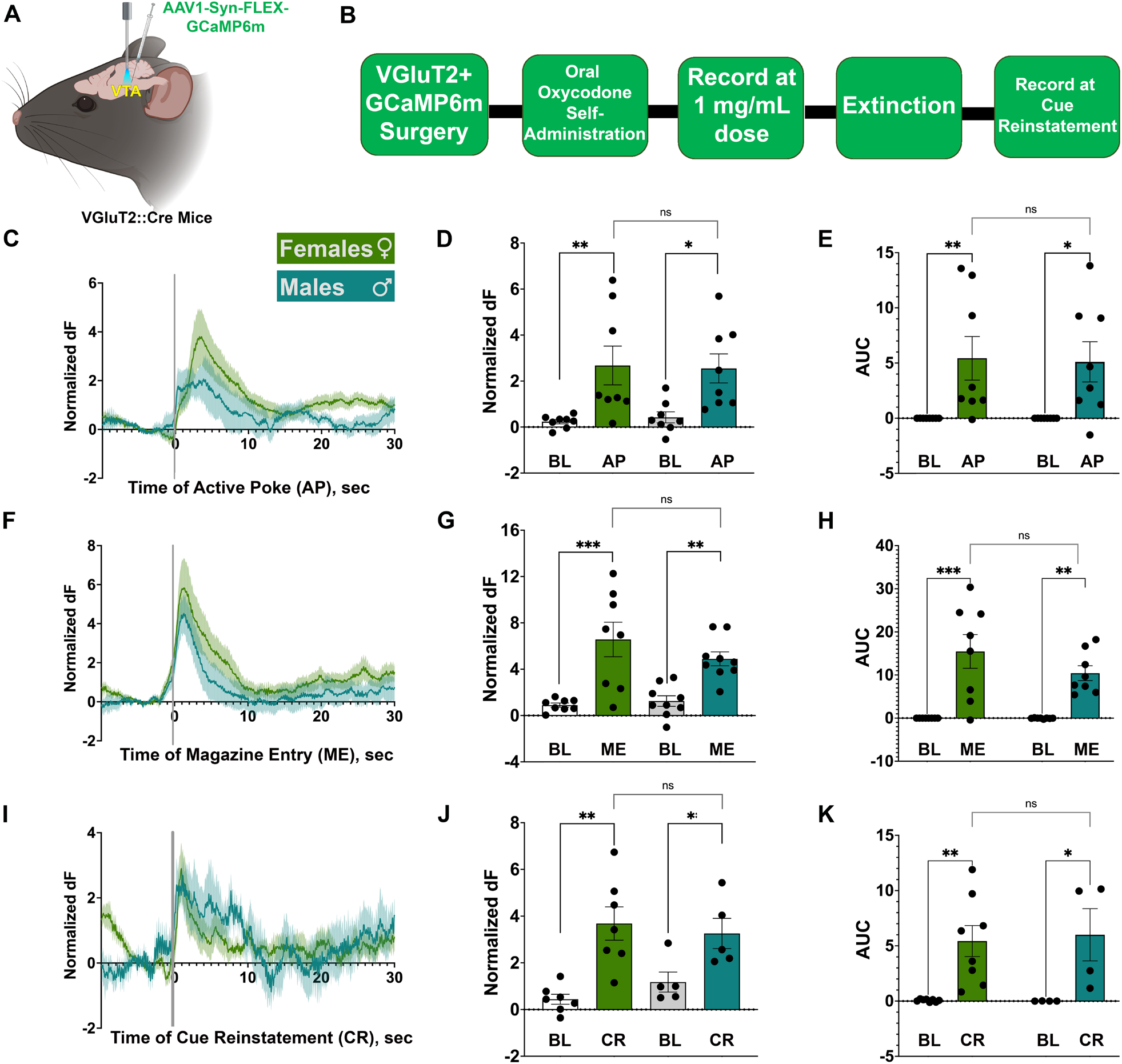
VTA VGluT2 neurons signal opioid self-administration-related behaviors. **A**. Surgical schematic: VGluT2-IRES::Cre male and female mice were injected in medial VTA with AAV-FLEX-GCaMP and were implanted with a fiber optic recording ferrule. **B**. Experimental timeline: 4 weeks following viral injection and implant, mice were trained on a FR2 reinforcement schedule and oxycodone dose was escalated every four days (0, 0.5, 0.1, 0.3, 0.5, 1.0 mg/mL). VGluT2 neuron calcium was recorded fiber photometrically at the highest dose. After reaching extinction criterion, mice were recorded during cued reinstatement. **C**. Normalized z-score for female (green) and male (blue) mice time locked to active poke (AP) for oxycodone reward during self-administration compared to baseline (BL). Mean represented by darker line and standard error represented by shaded line. **D**. Maximum values for individual mice following active poke (0-3 seconds poke) compared to baseline. Significant increases in both male and female signal following active poke compared to baseline but no significant sex difference. **E**. AUC analysis for recorded signals (0-3 seconds following poke). Significant increases in both male and female signal for active poke compared to baseline but no significant sex difference. **F**. Normalized z-score of both sexes following oxycodone magazine entry (ME) where consumption occurs. **G**. Maximum values for individual mice following magazine entry (0-3 seconds). Significant increases in both male and female signal compared to baseline but no significant sex difference. **H**. AUC analysis for recorded signals (0-3 following entry). Significant increases in both male and female signal for active poke compared to baseline but no significant sex difference. **I**. Normalized z-score of both sexes following cue presentation during cued reinstatement test (CR). **J**. Maximum values for individual mice following cue presentation (0-3 seconds). **K**. AUC for individual mice following cue presentation during reinstatement (0-3 seconds). Significant increases in both male and female signal for active poke compared to baseline but no significant sex difference.

After extinction of active responses in the absence of the drug-associated cue, active responses under extinction conditions resulted in the presentation of the drug-paired tone cue. VTA VGluT2+ calcium activity significantly increased following response-contingent presentation of the drug-cue, F(1,10) = 26.30, p=0.0004, for female mice: t(7)=4.749, p= 0.0016, and male mice: t(5)=2.702, p=0.044. Cue reinstatement response AUC was also significantly different from baseline, F(1,10)=20.33, p=0.0011, female mice: t(7)=3.634, p=0.0091 and male mice: t(5)=2.832, p=0.0353. There were no significant differences between sexes for maximum response F(1,10)=0.134, p=0.72 or AUC F(1,10)=0.032, p=0.86 (**Figure 4I-K**).

### VTA VGluT2 neuronal activity is required for cue-induced reinstatement of opioid-seeking behavior in male mice

Given that VTA VGluT2 GCaMP signal was increased by opioid-seeking behavior, we expressed the inhibitory designer receptor hM4Di or a fluorophore control in VTA VGluT2 neurons. We delivered a behaviorally subthreshold dose of clozapine[27], which we previously showed significantly reduces hM4Di-VGluT2 neuronal activity[25], during oxycodone self-administration or cue-induced reinstatement (**Figure 5A-B**). In male mice, chemogenetic inhibition of VTA VGluT2 neurons did not modify oxycodone consumption F(2,63)=1.218, p=0.3020 (**Figure 5C**). However, inhibition of VTA VGluT2 neurons significantly reduced active responding during cue-induced reinstatement following extinction of opioid-seeking behaviors, F(1,52)=8.437, p=0.0054 (**Figure 5D**). Inactive pokes were not changed by VGluT2 neuron inhibition, F(1,52)=0.5148, p > 0.05. In female mice, VTA VGluT2 neuron inhibition did not significantly alter oxycodone consumption F(2,30)=0.5864, p=0.5626 or reinstatement-related active responding, F(1,36) = 2.684, p=0.1101. Additionally, inactive poke responding was not altered by chemogenetic inhibition in female mice, F(1,37)=0.8589, p=0.3601 (**Figure 5E-G**). An additional cohort of male and female mice received identical DREADD or control surgeries and trained to orally self-administer water vehicle as a control for thirst behavior. DREADD inhibition of VTA VGluT2 neurons did not significantly alter water self-administration, F(1,28)=0.05567, p=0.8152, or cue reactivity in male or female mice following extinction, F(1,28)=0.05420, p=0.8176 (**Supplemental Figure 4**), suggesting that VTA VGluT2 neurons participate in oxycodone-seeking behavior rather than water-seeking behavior. To test the possibility that the reduced reinstatement of male mice following chemogenetic inhibition of VTA VGluT2 neurons resulted from changes in locomotor behavior, mice were injected with clozapine, placed in an open field and tracked for the following locomotor parameters: average speed, distance traveled, and maximum speed. There were no differences between any experimental condition for distance traveled F(3,56)=0.6438, p=0.5901, average speed F(3,56)=0.6488, p=0.5870, or maximum speed demonstrating F(3,56)= 1.152, p=0.3362. Thus, reduced reinstatement of oxycodone-seeking behavior in male mice was not driven by changes in locomotor behavior (**Supplemental Figure 4**).

**Figure 5.**
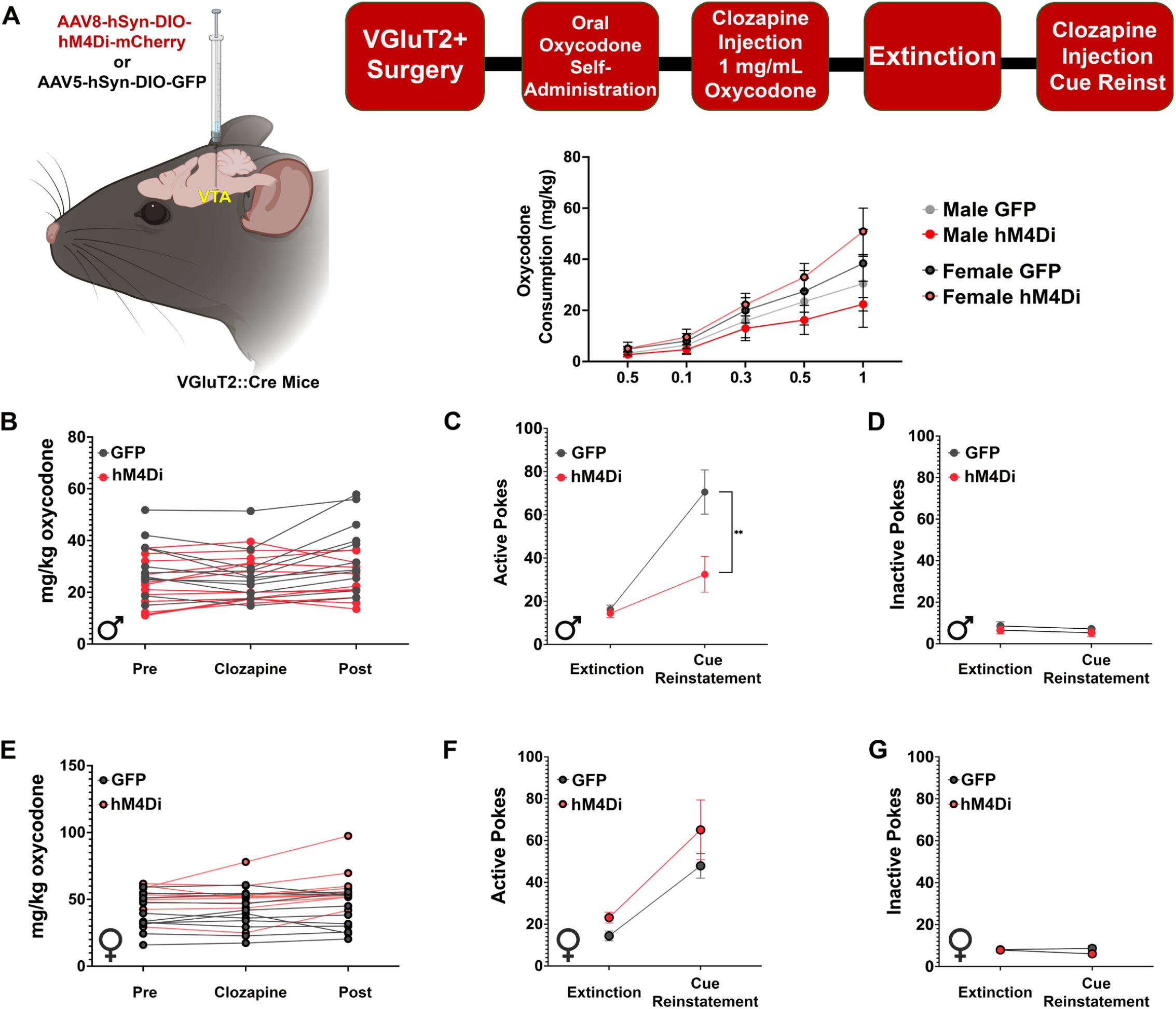
VTA VGluT2 neuron chemogenetic inhibition blocks oxycodone reinstatement in male but not female mice. **A**. Surgical schematic: VGluT2-IRES::Cre male and female mice were injected with AAV-FLEX-hM4Di or AAV-FLEX-GFP in the medial VTA. Experimental timeline: 4 weeks after viral expression, mice trained to orally administer drug on a FR2 reinforcement schedule and oxycodone dose was escalated every four days (0, 0.5, 0.1, 0.3, 0.5, 1.0 mg/mL). Behaviorally-subthreshold clozapine (0.1 mg/kg) was administered for one day at the highest dose, followed by 3 more days of self-administration. Mice were next trained to extinguish active responses and after extinction criterion was met, were administered clozapine (IP) during cued reinstatement testing. Subpanel: male and female mice differentially consumed oxycodone. **B**. Mg/kg oxycodone consumption for males, pre clozapine (3-day average), on clozapine day and post clozapine (3-day average). No significant changes in consumption for hM4Di group or GFP fluorescent control group. **C**. Recorded active pokes during cued reinstatement testing. VTA VGluT2 chemogenetic inhibition significantly reduced reinstatement behavior compared to GFP control group. **D**. Inactive pokes did not differ by experimental condition. **E**. Mg/mg oxycodone consumption for females. No significant changes in oxycodone consumption for hM4Di group or GFP fluorescent controls. **F**. Recorded active pokes during cue reinstatement testing. hM4Di inhibition did not alter reinstatement behavior in female mice. **G**. hM4Di inhibition did not alter inactive poke behavior.

## DISCUSSION

Understanding the circuit level neural contributors to opioid self-administration and relapse behavior are foundational to both treating opioid use disorder as well as developing efficacious pharmaceuticals with reduced addiction profiles. Mu opioid receptors (MORs) within the VTA are required for the reinforcing effects of opioids[6], are sufficient to induce reinstatement of extinguished drug-seeking behavior[33], and are canonically expressed in nondopaminergic GABA-releasing neurons[8]. Due to the cellular heterogeneity of VTA GABA neurons[28], we used *in situ* hybridization to determine the distribution of VTA VGluT2+VGaT―, VGluT2 +VGaT+, and VGluT2― VGaT+ neurons that express the MOR. The different types of VTA MOR-expressing neurons had distinct topographies across the anteroposterior VTA.

We found that most anterior VTA MOR-expressing neurons express VGluT2 without VGaT while most posterior VTA MOR-expressing neurons express VGaT without VGluT2. These data may be useful in determining why VTA opioid function differs across the anterior-posterior axis. For instance, while activation of the MOR is sufficient to induce reward selectively within the posterior VTA[34,35], AMPA receptors within the anterior VTA, where glutamate-releasing neurons are most dense[17,36] are selectively necessary for morphine reward[37]. In addition, overexpression of the AMPA receptor in anterior VTA enhances morphine reward while overexpression in the posterior VTA causes morphine to be interpreted as an aversive stimulus[38].

Based on the local GABA-MOR projection to VTA dopamine neurons, we hypothesized that VTA VGluT2 neuron projections to VTA dopamine neurons are also regulated by the MOR. Optogenetic stimulation of VTA VGluT2 neurons resulted in monosynaptic excitatory currents in VTA dopamine neurons, consistent with prior results[29,30,39]. Activation of MOR via bath application of DAMGO depressed the amplitude of VGluT2-CoChR evoked EPSCs in TH-expressing neurons, indicating that MOR modulates glutamate transmission to VTA dopamine neurons. The depression of VGluT2-CoChR evoked EPSCs with MOR activation may have been due to post-synaptic receptor desensitization[40]. However, activation of MOR via bath application of DAMGO increased the paired-pulse ratio. By demonstrating that MOR activation increases glutamate release probability in a paired-pulse stimulation, MOR is altering glutamate release by a presynaptic mechanism[41]. Therefore, our results indicate that the MOR is expressed on presynaptic terminals of VTA VGluT2 projections to VTA dopamine neurons. Though we identified all biocytin-labeled neurons from recordings as expressing tyrosine hydroxylase, we were unable to identify all recorded neurons and thus cannot exclude the possibility that non-dopamine neurons may also receive local MOR-gated inputs from VTA VGluT2+ neurons. Moreover, the local VGluT2+MOR+ projection identified here may be a subset of local projections from VGluT2+ neurons[29].

Having identified a local VTA VGluT2+MOR+ microcircuit to VTA dopamine neurons, we hypothesized that this pathway belonged to a macrocircuit resulting in dopamine release within the nucleus accumbens.

MOR agonists, such as heroin and morphine, cause the release of dopamine preferentially in the nucleus accumbens medial shell compared with the core[42,43]. Consistent with this, evidence suggests that VTA GABA-MOR neuron inhibition by heroin preferentially disinhibits VTA dopamine neurons that project to the medial shell[44]. A subset of VTA VGluT2 neurons co-releases dopamine to the nucleus accumbens medial shell[45]. We biased our GRABDA recordings to the nucleus accumbens core, which does not receive projections from VTA VGluT2+TH+ neurons[46-48], and thus dopamine released within the core results from the activation of VGluT2―TH+ neurons. We found that optogenetic activation of VTA VGluT2 neurons results in dopamine release within the accumbens core and that the magnitude of this response is reduced by the MOR agonist oxycodone *in vivo*. Thus, our results suggest that VTA VGluT2+MOR+ neurons regulate VTA dopamine neurons projecting to the accumbens core. Given that accumbal dopamine release is further regulated at release sites by multiple parallel circuits, further research will be necessary to test whether MOR-gated dopamine release resulting from a VTA VGluT2 neuron micro-circuit is regulated by cholinergic or other accumbal interneurons[49-53]. Further research will also be necessary to determine whether dopamine release within the nucleus accumbens medial shell, driven from VTA VGluT2+TH+ or VGluT2―TH+ neurons, is gated by VTA VGluT2+MOR+ neurons.

Nevertheless, because MOR activation decreased the excitatory post-synaptic current elicited by VTA VGluT2+MOR+ stimulation and subsequent dopamine release to the accumbens core, it is unlikely that VTA VGluT2+MOR+ neurons play a role in glutamatergic activation of VTA dopamine via the MOR. In other words, a different population of glutamate neurons is likely responsible for the glutamatergic activation of VTA dopamine neurons that follows MOR-induced GABA disinhibition [21].

In our investigation of a VTA VGluT2 neuron role in oral oxycodone self-administration and cue-induced reinstatement, we found that VTA VGluT2 neurons are significantly activated by different elements of self-administration and reinstatement. Population changes in VTA VGluT2 neuron calcium significantly increased following actions to acquire and consume oxycodone during self-administration and abstinent drug-seeking during reinstatement. Prior research has shown that subsets of VTA VGluT2 neurons are activated by reward-seeking actions, reward consumption, and reward-related cues[20,54]. Because the drug-associated cue was delivered following nose poke responses to acquire drug, further research will be necessary to disentangle whether the drug-seeking action, the drug-associated cue, or both, activate the same or different VTA VGluT2 neuron subtypes.

We have previously found that male mice consume less oxycodone (mg/kg) than female mice at the highest doses of self-administration[26]. We replicated this result and separated neuronal recording data between males and females to critically evaluate the potential contribution of VTA VGluT2 neurons to this behavioral difference. Neither phasic (maximum z-score change) nor tonic (AUC) measures of activity following any analyzed event significantly differed between males or female mice. However, during cue-induced reinstatement, chemogenetic inhibition of VTA VGluT2 neurons significantly reduced reinstatement behavior for male but not female mice. These findings demonstrate that VTA VGluT2 neurons play a causal role in abstinent cue-induced drug-seeking (relapse) behavior in at least male mice. It is possible that despite no identified sex differences in MOR distribution or VGluT2 neuronal signaling of drug-seeking behavior that circuit level differences exist between males and female mice that contribute to this discrepancy. Relatedly, we have found divergent roles of sex hormones between sexes that contribute to oxycodone-related behavior in mice[26]. Afferent circuitry to VTA VGluT2 neurons may also provide insight into this discrepancy and is of interest for future investigation.

For male and female mice, inhibition of VTA VGluT2 neurons had no effect on self-administration behavior. One reason for this may be that the presence of oxycodone during self-administration already inhibited VTA VGluT2+MOR+ neurons and thus chemogenetic inhibition would be mitigated. It is also plausible that disinhibition of VTA dopamine neurons resulting from activation of the MOR on midbrain GABA neurons and terminals overshadowed the effects of chemogenetic inhibition of VTA glutamate neurons during oxycodone self-administration.

Canonically, the reinforcing effects of opioids are thought to result from opioid activation of the MOR on midbrain GABAergic somata or terminals that disinhibit VTA dopamine neurons[8]. This two-neuron model of opioid addiction has been challenged by findings demonstrating that: 1) half of MOR-expressing axon terminals synapsing on TH-expressing dendrites are asymmetric (putative excitatory) or symmetric (putative inhibitory)[55]; 2) select subsets, rather than all VTA GABA or glutamate neurons, express the MOR[18,19]; 3) glutamate transmission within the VTA plays an essential role in the disinhibition of VTA dopamine neurons following MOR activation[21] as well as heroin self-administration[22]; and 4) ablation of the MOR within central and peripheral VGluT2 neurons results in reduced oxycodone self-administration, place preference, locomotion, and withdrawal[56]. Our results extend these findings by demonstrating that a subset of VTA VGluT2 neurons that lack VGaT expresses the MOR and participates in a local circuit that drives MOR-gated accumbal dopamine release.

We interpret these results to reflect that VTA dopamine neurons are regulated in a three-neuron model whereby 1) MOR activation results in increased mesolimbic dopamine release via disinhibition of VGaT+MOR+ neurons and 2) this mesolimbic dopamine release is restrained by MOR-dependent depression of excitatory synaptic drive from local VTA VGluT2+MOR+ inputs onto accumbens-projecting VTA dopamine neurons. Due to the dense interconnected nature of VTA neurons, an additional MOR sensitive microcircuit between VGluT2+MOR+ and VGaT+MOR+ local populations may also play a regulatory role in subsequent dopamine release but has yet to be explored. Further research will be needed to identify if the same or different VTA dopamine neurons are regulated by VTA VGaT+MOR+ or VGluT2+MOR+ neurons. However, because heroin preferentially activates VTA dopamine neurons projecting to the nucleus accumbens medial shell[44], and we detected MOR-regulated dopamine release in the nucleus accumbens core as a result of activating VTA VGluT2 neurons, we hypothesize that each VTA MOR+ cell-type regulates distinct accumbal dopamine release targets to differentially regulate self-administration versus abstinent drug-seeking (relapse/reinstatement). In this model, 1) reduction of VGaT +MOR+ neurons by MOR activation disinhibits shell-projecting dopamine neurons to drive opioid reinforcement; and 2) because D1 receptors in accumbens core, but not shell, are required for cue-induced reinstatement of opioid seeking[57], activation of VGluT2+MOR+ neurons results in the activation of core-projecting dopamine neurons to drive abstinent drug seeking.

## MATERIALS AND METHODS

### Animals

Male and female C57BL/6J and VGluT2-IRES::Cre mice (Slc17a6tm2(cre)Lowl/J; Stock #016963) were purchased from The Jackson Laboratory (Bar Harbor, ME) and bred at the University of Colorado. Mice were maintained in a colony room with a 12-hr light/dark cycle, and and water ad libitum. Mice were between 8-20 weeks at the time of surgical procedures. All animal procedures were performed in accordance with the National Institutes of Health Guide for the Care and Use of Laboratory Animals and approved by the University of Colorado Boulder Institutional Animal Care and Use Committee. *Ex vivo* electrophysiological experiments were approved by George Washington University Institutional Animal Care and Use Committee.

### In situ hybridization and analysis of VTA cell-types

Brains were rapidly extracted from male (n=4) and female (n=3) C57BL6J mice (8-12 weeks old). VTA sections were mounted onto Fisher SuperFrost Plus slides and RNAscope *in situ* hybridization was performed according to the manufacturer’s instructions. Briefly, sections were treated with heat and protease digestion followed by hybridization with target probes to mouse Oprm1 (ACDBio #489311), VGluT2 (ACDBio #319171), and VGaT (ACDBio #319191). Additional sections were hybridized with the bacterial gene DapB as a negative control, which did not exhibit fluorescent labeling. Slides were coverslipped in Prolong Diamond with DAPI (ThermoFisher) and imaged on a Nikon A1R confocal (20X) with 3 µm steps z-stacked images at 512 × 512 pixel resolution. For cell counts, each section was counted by three independent scorers. Only fluorescence with DAPI co-labeling was counted. A minimum of four transcripts was required for classification of one cell. To analyze differences in anteroposterior cell-types, the VTA was split equally in four sections each by the full emergence of the interpeduncular nucleus: anterior -3.08 to -3.40 and posterior -3.52 to -3.88 mm from bregma.

### Stereotactic surgery

Male and female VGluT2-IRES::Cre mice were anesthetized with 1-3% Isoflurane and secured in a stereotactic frame (Kopf). AAV8-hSyn-DIO-hM4Di-mCherry (Addgene, Cambridge, MA), AAV5-hSyn-DIO-GFP (Addgene), or AAV1-Syn-FLEX-GCaMP6m (Addgene), or AAV8-hSyn-DIO-hM4Di-mCherry (Addgene), AAV8-hSyn-FLEX-CoChR-GFP (UNC Vector Core), AAV8-nEF-Con/Foff-ChRmine-oScarlet (Addgene) was injected into VTA (AP: -3.2 mm; ML: 0.0 mm; DV: -4.3 mm). All AAVs were injected at 2-5 × 10^12 titer. Injection volume (500 nL) and flow rate (100 nL/min) were controlled with a microinjection pump (Micro4; World Precision Instruments, Sarasota, FL). Following injection, the needle was left in place for an additional 10 min for virus diffusion, after which the needle was slowly withdrawn. Nucleus accumbens surgeries followed the same protocol with the modification of virus: AAV9-hSyn-GRABDA1h (coordinates AP: 1.34, ML: 1.1 DV: -4.5). For GCaMP recordings, mice were additionally implanted with an optic fiber (400 μm core diameter, 0.66 NA; Doric Lenses, Quebec, Canada) dorsal to VTA (AP: -3.2 mm relative to bregma; ML: -1.0 mm at 9.5°; DV: -4.2 mm) and for GRABDA recordings implanted dorsal to nucleus accumbens (AP: 1.34, ML: 1.1 DV: -4.3) that was secured with screws and dental cement to the skull. For ChRmine stimulation, mice were implanted with an optic fiber (200 μm core diameter, Doric) dorsal to VTA (AP: -3.2 mm; ML: -1.0 mm at 9.5°; DV: -4.2 mm) that was secured with screws and dental cement to the skull. All mice were allowed to recover for 3-4 weeks before experimentation

### Acute slice preparation

Male (n=4) and female (n=3) VGluT2-IRES::Cre mice were injected in VTA with AAV8-hSyn-FLEX-CoChR-GFP (UNC Vector Core, 500 nL, 100 nL/min, 4 x10^12 titer). After one week of acclimation, acute coronal slices were prepared as previously described [23] from mice deeply anesthetized with a mixture of ketamine (100 mg/kg) and dexmeditomidine (0.25 mg/kg). Mice were perfused with 34°C NMDG ringer: 92 mm NMDG, 2.5 mm KCl, 1.2 mm NaH2PO4, 30 mm NaHCO3, 20 mm HEPES, 25 mm glucose, 5 mm sodium ascorbate, 2 mm thiourea, 3 mm sodium pyruvate, 10 mm MgSO4, and 0.5 mm CaCl2[24] Following perfusion, the brain was rapidly dissected and horizontal slices (220 μm) were prepared in warmed NMDG ringer using a vibratome. Slices recovered for 1 h at 34°C in oxygenated HEPES holding solution: 86 mm NaCl, 2.5 mm KCl, 1.2 mm NaH2PO4, 35 mm NaHCO3, 20 mm HEPES, 25 mm glucose, 5 mm sodium ascorbate, 2 mm thiourea, 3 mm sodium pyruvate, 1 mm MgSO4, and 2 mm CaCl2 [24] and then were held in the same solution at room temperature until use.

### Slice electrophysiology

For electrophysiological recordings, a single slice was transferred to a chamber perfused at a rate of 1.5 to 2.0 ml/min with heated (28-32C) artificial cerebrospinal fluid (aCSF, in mM: 125 NaCl, 2.5 KCl, 1.25 NaH2PO4, 1 MgCl2 6H2O, 11 glucose, 26 NaHCO3, 2.4 CaCl2, saturated with 95% O2 and 5% CO2. 100 µM picrotoxin was included in all recordings to pharmacologically isolate EPSCs. Non-GFP expressing cells were visually identified and selected for recordings. Whole-cell patch-clamp recordings were performed using a Sutter IPA amplifier (1 kHz low-pass Bessel filter and 10 kHz digitization) using SutterPatch software (Sutter Instruments). Voltage-clamp recordings at -70 mV were made using glass patch pipettes with resistance 2-4 MOhms, filled with internal solution containing (in mM):. 0.4% biocytin was included in the internal solution for all recordings. Series resistance was monitored throughout voltage clamp recordings and recordings in which the series resistance changed more than 20% and/or exceed 20 MOhms were not included in the analysis. Junction potential was not corrected for.

EPSCs were optically evoked using an optical fiber (Plexon, Inc, Dallas, TX) coupled to an 465 nm LED and driver (Plexon, Inc, Dallas, TX). Light pulses (0.5-4 ms) were triggered by TTL pulses from the SutterPatch software. EPSCs were stimulated every 30 seconds to avoid desensitization of CoChR. After a stable 10 minute baseline of EPSCs was collected, DAMGO (500 nM, Bio-Techne, Minneapolis, MN) was bath applied for 10 minutes. Data was analyzed offline using SutterPatch software and Graphpad Prism 9.0. To create time-courses, EPSC amplitudes were binned in 2-minute bins and normalized to the last 5 minutes of the baseline. To determine average amplitudes pre- and post-DAMGO, EPSC amplitudes were averaged over the last 5 minutes of the baseline and the last 5 minutes of DAMGO application. Representative traces are an average of 10 consecutive sweeps immediately proceeding application or removal of DAMGO.

### In vivo stimulation and optical dopamine recordings

Male and female VGluT2-IRES::Cre mice were injected in VTA with AAV8-nEF-Con/Foff-ChRmine-oScarlet (500 nL, 100 nL/min, 5 x10^12 titer) and in nucleus accumbens AAV9-hSyn-GRABDA1h (500 nL, 100 nL/min, 5 × 10^12). Animals were tail restricted to minimize potential movement-related GRABDA signaling. Five minutes prior to stimulation and recording, mice received an IP injection of saline (pre-test), oxycodone (10 mg/kg), or saline (post-test). Mice received 5 stimulation trials for each frequency (5,10, 20, 40 Hz). Order was counterbalanced and randomized for each mouse per experimental condition. Total stimulation time (2.5 sec per train) was controlled per frequency and mice were given a 5 minute break between each stimulation trial for signal to return to baseline. Stimulation parameters per frequency: 5hz (5ms width, period 200ms, count 13) 10hz (5ms width, period 100ms, count 25) 20hz (5ms width, period 50ms, count 50) 40hz (5ms width, period 25ms, count 100). Mice were given three days in between each injection. GRABDA was excited at two wavelengths (465 nm and 405 nm control) with amplitude-modulated signals from two light-emitting diodes reflected off dichroic mirrors and then coupled into an optic fiber[25]. Signals from GRABDA and the control channel were returned through the same optic fiber and acquired with a femtowatt photo-receiver (Newport, Irvine, CA), digitized at 1kHz, and then recorded by a real-time signal processor (TDT). Timestamps for optical stimulation were digitized in Synapse software (TDT). Analysis of the recorded GRABDA signal was performed using custom-written MATLAB scripts. Signals (465 nm and 405 nm) were downsampled (10X) and peri-event time histograms (PETHs) were created trial-by-trial between -10 sec and +30 sec surrounding each optical train onset. For each trial, data was detrended by regressing the control signal (405 nm) on the GRABDA signal (465 nm) and then generating a predicted 405 nm signal using the linear model generated during the regression. The predicted 405 nm channel was subtracted from the 465 nm signal to remove photo-bleaching and fiber bending artifacts (ΔF).

### Self-administration

Training commenced as previously described[26]. Briefly, mice trained on a FR1 schedule of reinforcement for water over three days and an FR2 schedule of reinforcement for water over three additional days. Within the active port, a cue light was illuminated until reward was earned. At reward, the cue light was terminated, a 7 kHz tone was sounded together over 10 sec, and 20 µL reward was delivered. After 10 sec, the cue light was re-illuminated. For oxycodone mice, FR2 oxycodone administration followed an escalating dose schedule (0.05, 0.1, 0.3, 0.5, 1.0 mg/mL) for three days per dose and 3 hour sessions per day under the same reward delivery conditions. For water mice, the same number of self-administration sessions were made as oxycodone mice, but the reward was always water vehicle. After three days at the highest dose, or the matched number of training days for water mice, all mice began extinction where active and inactive nose pokes were counted but had no programmed consequence. Extinction sessions were 1 hour and criterion for extinction was calculated as 30% of the average total active pokes for each mouse at their highest dose. Mice were required to emit less than or equal to this active poke criterion for two consecutive days before shifting to cued reinstatement testing. The following day after reaching extinction criterion, single active nose pokes activated the cue previously associated with oxycodone reward consisting of a 10-second 7 kHz tone together with the termination of a light within the active port. Inactive nose pokes were counted without other programmed consequence. Mice remained in the cue phase for 1 hour following the first earned cue presentation.

### GCaMP Recordings

Male and female VGluT2-IRES::Cre mice were injected in VTA with AAV1-Syn-FLEX-GCaMP6m four weeks prior to behavioral testing. At least four weeks following intra-VTA viral injection, GCaMP6m recordings were performed in while mice operantly earned oxycodone (1 mg/mL) and during cued reinstatement testing. GCaMP6m was excited at two wavelengths (465 nm and 405 nm isosbestic control) with amplitude-modulated signals from two light-emitting diodes reflected off dichroic mirrors and then coupled into an optic fiber[57]. Signals from GCaMP and the isosbestic control channel were returned through the same optic fiber and acquired with a femtowatt photo-receiver (Newport, Irvine, CA), digitized at 1kHz, and then recorded by a real-time signal processor (Tucker Davis Technologies). Behavioral timestamps for active pokes, inactive pokes. magazine entry, and cue presentation were digitized in Synapse by TTL input from MED-Associates.

Analysis of the recorded calcium signal was performed using custom-written MATLAB scripts. Signals (465 nm and 405 nm) were downsampled (10X) and peri-event time histograms (PETHs) were created trial-by-trial between -10 sec and +30 sec surrounding each event. For each trial, data was detrended by regressing the isosbestic control signal (405 nm) on the GCaMP signal (465 nm) and then generating a predicted 405 nm signal using the linear model generated during the regression. The predicted 405 nm channel was subtracted from the 465 nm signal to remove movement, photo-bleaching, and fiber bending artifacts (ΔF). Baseline normalized maximum z-scores were taken from -3 to 0 seconds prior to programmed time stamps of interest (active/inactive poke, cue, reward entry) and maximum shock z-scores were taken from 0 to 3 seconds following behavioral events. AUC analysis per animal were normalized to the baseline 3 seconds prior to shock and 3 seconds following programmed behavioral events.

### Chemogenetic inhibition of VTA glutamate neurons during oxycodone self-administration and reinstatement

Male and female VGluT2-IRES::Cre mice were injected in VTA with AAV8-hSyn-DIO-hM4Di-mCherry or AAV5-hSyn-DIO-GFP four weeks prior to behavioral testing, similar to prior experiments examining the roles of VTA VGluT2 neurons in motivated behavior[25]. Mice self-administered oxycodone or water as described. However, after three days of self-administration at the highest dose, mice received an IP injection of a behaviorally-subthreshold[27] dose of clozapine (0.1 mg/kg) ten minutes prior to self-administration. This dose has previously been shown to decrease VTA VGluT2 neuron activity[25]. After the clozapine session, three additional days of training at 1 mg/mL commenced to compare self-administration before and after chemogenetic manipulation of VTA VGluT2 neurons. After self-administration, active responding was extinguished as described and the following day after mice met extinction criterion, an IP subthreshold dose of clozapine (0.1 mg/kg) was given ten minutes prior to cued reinstatement testing. Two days after reinstatement testing mice were given IP injection of clozapine (0.1 mg/kg) and placed in an open field. AnyMaze tracking was used to evaluate distance traveled, maximum distance, and average speed per mouse.

### Histology

VGluT2-IRES::Cre mice were anesthetized with isoflurane and perfused transcardially with phosphate buffer followed by 4% (w/v) paraformaldehyde in 0.1 M phosphate buffer, pH 7.3. Brains were extracted, post-fixed overnight in the same fixative and cryoprotected in 18% sucrose in phosphate buffer at 4°C. Coronal sections containing the VTA (30 μm) were taken on a cryostat, mounted to gelatin-coated slides, and imaged for mCherry (tethered to hM4Di), GFP, or GCaMP6m, or GRABDA expression and optical fiber cannula placement on a Zeiss Axioscope depending on experimental condition. Mice with optic fibers not localized to recording VTA GCaMP-expressing neurons were removed from the study. Sections used for whole-cell recordings, washed in phosphate buffer, blocked in 4% bovine serum albumin supplemented with 0.3% Triton-X 100 for one hour, incubated over three days at 4°C with anti-tyrosine hydroxylase (MAB318, 1:100), followed by washing in phosphate buffer, and incubating over three days at 4°C with streptavidin-conjugated Alexa647 (Jackson Secondaries 016-600-084, 1:50) and anti-mouse Alexa594 (Jackson secondaries 715-585-150, 1:50). After washing in phosphate buffer, slides were coverslipped in FluoroMount-G (ThermoFisher) and imaged on a Nikon A1R confocal (20X). After visual inspection to locate the biocytin-filled Alexa647-labeled neuron, 3 µm step z-stack images were made at 512 × 512 pixel resolution.

### Statistics

Tests were conducted in SPSS (IBM) or Prism (GraphPad Software; San Diego, CA). GCaMP data was analyzed by comparing the maximum GCaMP z-score during baseline, 3 seconds prior to event compared to the maximum z-score following the following events 3 seconds following event initiation: active poke, magazine entry, and cue presentation during reinstatement using Wilcoxon sign ranked tests. Identical analysis was performed on AUC statistics from the same behavioral timepoints. Max GRABDA normalized z-score values were averaged across for all trials per frequency and experimental condition (saline versus oxycodone), with 3 days between each stimulation condition. For GRABDA recordings, maximum fluorescent baselines or stimulated peaks did not differ between male or female mice, therefore data was collapsed across sex. Repeated measured ANOVA compared fluorescent change across experimental conditions per frequency. A between-subjects t-test analyzed the difference active and inactive pokes during cued reinstatement. A repeated measures ANOVA was used to compare baseline oxycodone consumption to hM4Di test day consumption and to post-test baseline. Separate analyses were performed for each sex. For all ANOVAs, if the assumption of sphericity was not met (Mauchley’s test), the Greenhouse-Geisser correction was used and Sidak-adjusted pairwise comparisons followed up significant main effects or interactions. Slice electrophysiology data were analyzed using paired t-test.

## Acknowledgments

We thank Alysabeth Phillips for technical assistance.

## Author Contributions

D.J.W., A.M.P., and D.H.R. conceived and performed experiments, wrote the manuscript, and secured funding. C.M., A.L., E.D.P. performed experiments and contributed to the writing of the manuscript.

## SUPPLEMENTAL

**Supplemental Figure 1.**
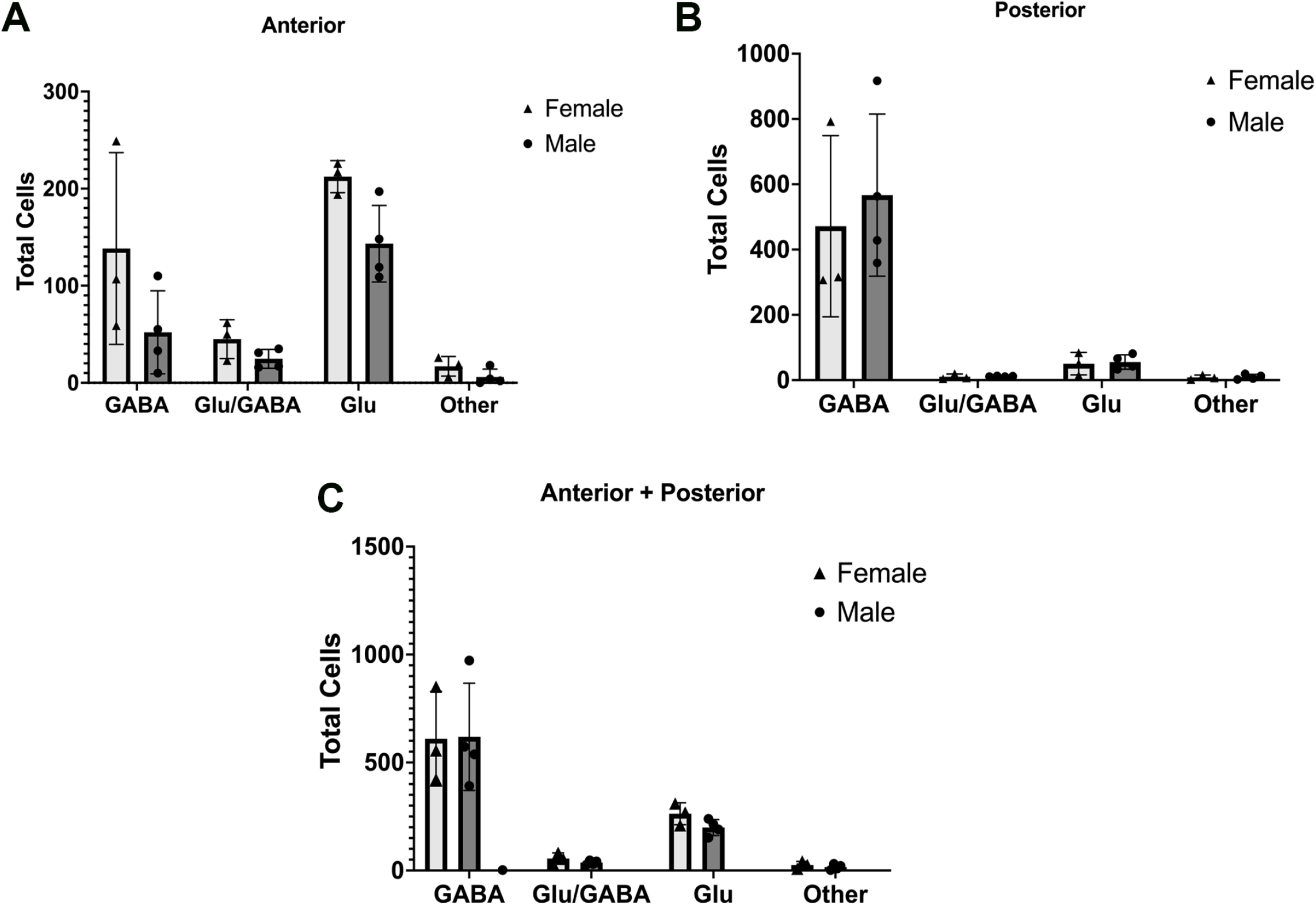
MOR distribution across the anteroposterior extent of the VTA. **A**. Anterior quantification of MOR+ cells corresponding to VGluT2 subtypes (GABA: VGluT2―VGaT+MOR+, Glu/GABA: VGluT2+VGAT +MOR+, Glu: VGluT2+VGaT―MOR+). **B**. Posterior VTA quantification of same cell-type categories. **C**. Collapsed data for both anterior and posterior VTA topography.

**Supplemental Figure 2.**
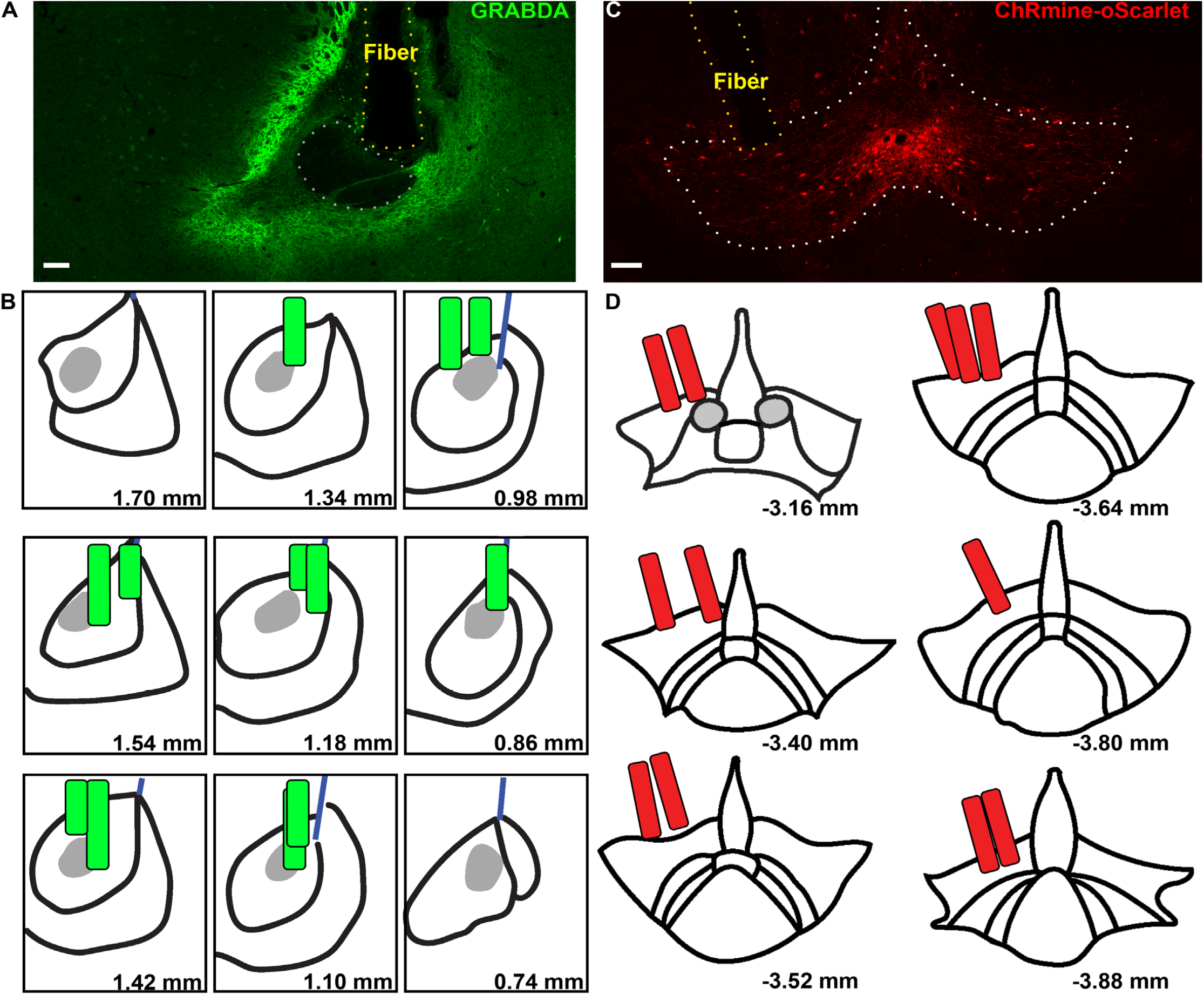
Localization of GRABDA recording fibers in nucleus accumbens and stimulating fibers in VTA VGluT2 ChRmine neurons. **A**. Example recording fiber (yellow outline) within nucleus accumbens (anterior commissure outlined in gray). Green fluorescence is GRABDA1h. **B**. Histological localization of recording fibers in nucleus accumbens based on Franklin and Paxinos1 numbers anterior to bregma. Gray is anterior commissure. Blue is lateral ventricle. **C**. Example stimulating fiber (yellow outline) within VTA (outlined in white). Red fluorescence is ChRmine-oScarlet. **D**. Histological localization of stimulating fibers in VTA based on Franklin and Paxinos1 numbers posterior to bregma. Gray is fasciculus retroflexus. Scale bars in A and C = 100 µm.

**Supplemental Figure 3.**
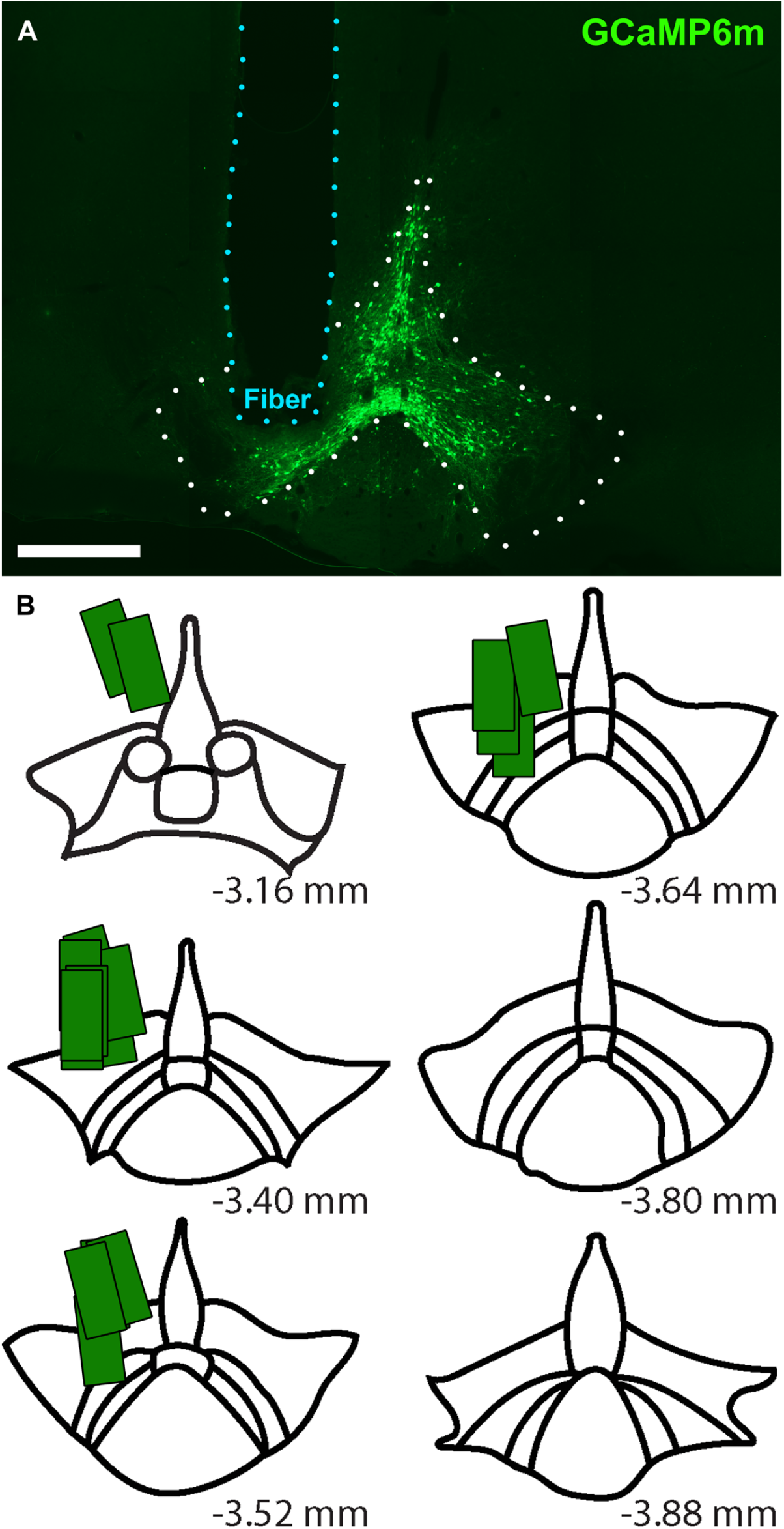
Localization of fibers recording VTA VGluT2 GCaMP6m signals. **A**. Example fiber (cyan outline) within VTA (outlined in white). Green = GCaMP6m neurons. Scale bar is 500 µm. **B**. Histological localization of all fibers recording VTA VGluT2 GCaMP6m neurons. Numbers refer to posterior position from bregma.

**Supplemental Figure 4.**
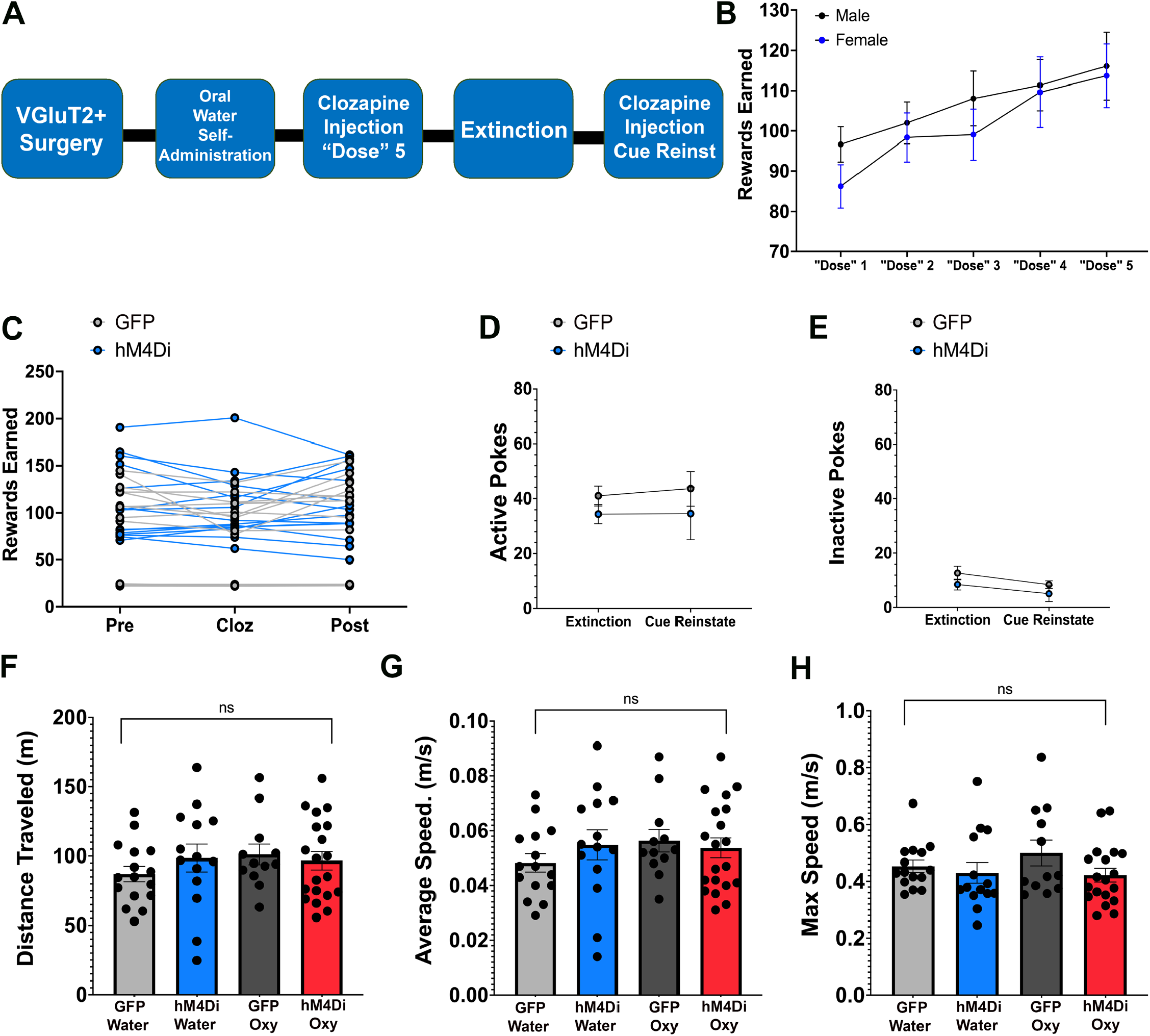
Water self-administration and locomotion behavior. A. Experimental timeline. **A**. VGluT2-IRES::Cre male and female mice received an AAV encoding FLEX-hM4Di or FLEX-GFP in the medial VTA. 4 weeks after viral expression, mice trained to orally self-administer water for the same number of sessions as oxycodone mice. Session averaging was matched to oxycodone group to establish “dose”. **B**. Rewards earned for water self-administration. Because no significant differences were observed between males and females, groups were collapsed. **C**. Rewards earned prior to clozapine (3-day average), on clozapine day, and post clozapine (3-day average). Chemogenetic VGluT2 neuron inhibition did not alter water self-administration. **D**. Active pokes during cued reinstatement testing. Mice did not significantly reinstate in response to water-paired cues and no significant differences in experimental groups following clozapine administration were observed. **E**. Inactive pokes were not altered by reinstatement or clozapine. **F-H**. No significant changes in distance traveled (m/s) (F) average speed (m) (G) or max speed (m/s) (H) between experimental groups following clozapine administration.

